# Natural variation in circadian period correlates with diverse phenological measures in *Boechera stricta*

**DOI:** 10.1101/2024.04.15.589576

**Authors:** Rob McMinn, Matti J. Salmela, Cynthia Weinig

## Abstract

The circadian clock is a time-keeping mechanism that enables adaptive responses to temporal environmental changes. Circadian period exhibits significant segregating genetic variation within and among populations along an elevational cline, potentially resulting from variable selection across microsites on outputs of the circadian clock. Reproductive timing is an important clock output and selection may favor delayed phenology. We performed a three-year common garden field study with the short-lived perennial *Boechera stricta* to quantify life history differences among 20 populations from an elevational gradient and among 20 maternal families within one population. We measured days to bolting and later life history transitions such as days to flowering; in the same genotypes, we measured circadian period. The timing of all life history transitions varied among growing seasons, suggesting adaptive life history evolution to local climate conditions. Life history transitions after bolting were also associated with circadian period, such that lengthened period was associated with delayed life history transitions. Structural equation modeling indicated that indirect selection via days to flowering and fruit production favored lengthened clock period in the low elevation common garden site, suggesting that selection on phenology could explain the evolution of variable circadian periods observed among populations from differing environments.

## Introduction

Natural populations experience temporal fluctuations in environmental conditions on multiple scales, necessitating sensitive time-keeping mechanisms. For instance, natural populations experience exogenous diel light−dark cycles on a 24-hour basis, variation in the length of a growing season, and seasonal changes within a year (Hut *et al*., 2013). The endogenous circadian clock is sensitive to several abiotic signals, including differences in photoperiod and temperature that presage temporal environmental changes (McClung, 2006; Greenham and McClung, 2015). Different endogenous timekeeping systems that are necessary for adaptation to fluctuating environments are likely to be interrelated at the genetic level: based on theoretical modeling, the molecular complexity of the plant circadian clock is indicative of responsiveness to both daily and seasonal environmental cycles (Troein *et al*., 2009), and studies on mutant genotypes, experimental mapping populations and natural accessions of *A. thaliana* have found that circadian clock timing regulates phenotypes from the molecular to organismal level over temporal scales of hours and days to seasons (Covington *et al*., 2008; Seo *et al*., 2012; Graf *et al*., 2017).

Given its pervasive phenotypic effects, circadian clock regulation could be an underlying component of plant adaptation to seasonal environments along latitudinal or elevational gradients. Variable selection on circadian clock outputs across environmental gradients could also explain the substantial segregating variation in circadian parameters within and among natural populations. In natural environments, clock cycles have a periodicity close to the day length of 24 hours. The classical circadian resonance hypothesis postulates that a functional clock with an endogenous period length close to 24 hours positively affects performance under 24-hour exogenous cycles (Pittendrigh and Minis, 1972; Ouyang *et al*., 1998; Woelfle *et al*., 2004; Dodd *et al*., 2005; Rubin *et al*., 2017), by allowing organisms to anticipate external changes and adaptively adjust their physiology (Sanchez and Kay, 2016). However, several common garden studies have shown significant natural genetic variation for circadian parameters segregating in wild populations that experience only 24-hour diurnal cycles in their habitats (Salmela and Weinig, 2019). For example, circadian period varied by 6.5 hours across a global collection of 150 *A. thaliana* genotypes (Michael *et al*., 2003). Segregating clock variation is also observed over smaller geographic scales. Salmela *et al*. (2016) and McMinn *et al*. (2022) sampled natural populations that consisted of multiple genotypes and found that quantitative genetic variation in circadian period segregated both among and within local populations of the North American Arabidopsis relative *Boechera stricta* found in the central Rocky Mountains. McMinn *et al*. (2022) also found that elevation explained a greater proportion (23%) of among-population variation in circadian period in a regional sample of *B. stricta* than did latitude in a global genotypic sample of *A. thaliana* (∼7%) (Michael *et al*., 2003), with shorter circadian periods and less within-population variation observed at higher elevations. Furthermore, the magnitude of intrapopulation variation was high, with genotypic ranges in circadian period of up to 5.8 hours on a spatial scale of only a few hundred meters (McMinn *et al*., 2022). Collectively, these findings suggest that beyond the classical circadian resonance hypothesis, other explanations, such as variable natural selection on phenotypic outputs, need to be scrutinized to understand the causes of naturally occurring genetic variation in the circadian clock.

The timing of life history events is a well-described output of the circadian clock and important in adaptation to spatial and temporal heterogeneity (Endler, 1986; Kawecki and Ebert, 2004). Many evolutionary studies have quantified the association between fitness and the developmental transition to reproduction (Stinson, 2004; Donohue *et al*., 2005; Ehrlén and Münzbergová, 2009). In strongly seasonal environments like those at high latitudes or elevations where the growing seasons are limited in duration, delayed flowering time may compromise fruit set and fitness; alternatively, delayed flowering may be advantageous in settings with longer growing seasons (Izawa, 2007; Leinonen *et al*., 2013; Yan *et al*., 2021), either because more meristems are produced when the period of vegetative growth is extended or because greater carbon resources accumulate (Geber, 1990). Multiple stages of life-history transitions are distinguishable, such as the transition from vegetative growth to reproduction (bolting and flowering), the transition to first fruiting, and the completion of all fruit production, although only the first of these has been extensively examined. Genetic variation in phenology traits is ubiquitous in a variety of species that range from short-lived and small model species like *Arabidopsis thaliana* (e.g., Méndez-Vigo *et al*., 2011; Vidigal *et al*., 2016) to long-lived and large forest trees (e.g., Mimura and Aitken, 2010; Alberto *et al*., 2013). Quantitative variation in these phenological traits arises due to segregating variation at loci acting in diverse developmental pathways, such as the circadian clock pathway, and many studies in model systems and crop species have indeed shown that the circadian clock affects flowering time (Schaffer *et al*., 1998; Brachi *et al*., 2010; Johansson and Staiger, 2015; Rubin *et al*., 2018; Rees *et al*., 2021). Importantly, in perennials, selection on optimal reproductive timing may be consistent or variable across years, depending on interannual environmental variation, and consistent selection over multiple growing seasons would be a prerequisite to allele frequency changes in the underlying pathways.

Joint examination of circadian phenotypes, life history traits, and components of plant fitness holds promise in deciphering the evolutionary forces that could shape natural variation in the circadian clock along various environmental gradients (Helm *et al*., 2017; Salmela and Weinig, 2019). Here, we explore potential evolutionary mechanisms contributing to the observed magnitude of natural genetic variation in circadian period by testing for potential covariation between circadian periodicity and phenology among populations living along an 800m elevational gradient in southeastern Wyoming, a region that encompasses marked environmental heterogeneity and includes natural populations of *B. stricta* that exhibit significant genetic differentiation in circadian period (a 3h range among population means (McMinn *et al*., 2022)). Due to strongly elevation-dependent variation in temperature and precipitation that limit growing season length, regional genetic variation in circadian traits might be maintained via natural selection on phenological traits that facilitate adaptation to highly seasonal environments. In a common garden in a field setting, we measured phenological and size traits over the course of three years for individuals from 20 populations derived from an elevational span of 800 meters and from 20 maternal genotypes derived from a single population. Using a multi-year common garden study, our research addresses *1)* how genetic variation for phenology and size is partitioned in multiple growing seasons among and within populations derived from an elevational gradient, *2)* whether the expression of phenological traits correlates with circadian period, and *3)* whether circadian period is directly or indirectly associated with measures of reproductive fitness. We hypothesize that directional selection on plant phenology has resulted in higher-elevation populations from sites with shorter growing seasons evolving earlier phenology relative to those from lower elevations. Further, should common genes regulate both the circadian clock and phenological traits, we anticipate significant correlations to arise between circadian period and reproductive phenology both among and within populations, such that shorter circadian periods (or the correlated expression of advanced circadian phase) are associated with early phenology (McClung, 2006) as is the case in molecular genetic studies of circadian clock regulation. Finally, to elicit evolutionary changes in circadian clock timing, phenology selection must be consistent over multiple growing seasons.

## Methods

### 2.1 Study species

*Boechera stricta* (Graham) Al-Shehbaz is a short-lived perennial plant and a member of the Brassicaceae family and has become an important model species for evolutionary ecology (Rushworth *et al*., 2011; Rushworth *et al*., 2022). The species is predominantly self-fertilizing (Song *et al*., 2006), and populations of the plant are distributed throughout North America, in particular through the Rocky Mountain region of the western United States. The species has been studied for local adaptation to diverse settings (Lee and Mitchell-Olds, 2013; Lin *et al*., 2021), for mechanisms acting in the maintenance of genetic variation (Anderson *et al*., 2013), and as a model to study the genetic architecture of flowering (Yan *et al*., 2021).

### 2.2 Common garden experiment

To partition variation in the expression of life history traits within *vs*. among populations and to test if life-history variation correlates with segregating variation in the circadian clock, we designed a common garden experiment (Clausen and Hiesey, 1958). The experiment included 20 geographically well-separated populations and 20 maternal families from within one population (Figure 1). Seeds from 20 natural populations were collected in 2015 across an elevational and longitudinal gradient from the northern extent of the Rocky Mountains, including the Medicine Bow, Sierra Madre, and Laramie Ranges. The gradients spanned 2500m to 3150m in elevation and 150km from eastern to western sites (Figure 1). For the common garden experiment, we selected 1) 50 replicates representing 10 maternal families that were pooled together and come from each of 20 populations (N = 1000 total replicates) and 2) 50 replicates of 20 maternal family lines from the Barber Lake population (denoted as BBL in Figure 1; 2625m elevation, 41.323 degrees N, 106.176 degrees W) (N = 1000 total replicates). The genetic structure among the experimental plants was chosen to enable comparison of within vs among-population variation, and the BBL population was chosen for its central position in the distribution of experimental populations. Seeds were planted into pots filled with Redi-Earth potting mix (Sungro, Agawam, MA, USA) and placed in a greenhouse at the Agricultural Experiment Station (Laramie, WY) to germinate. Seedlings were grown in the greenhouse for six weeks, at which time the majority of plants had four true leaves. Seedlings were then transplanted directly into the soil in a field plot outside the University of Wyoming Agricultural Experiment Station in the second week of October.

**Figure 1.**
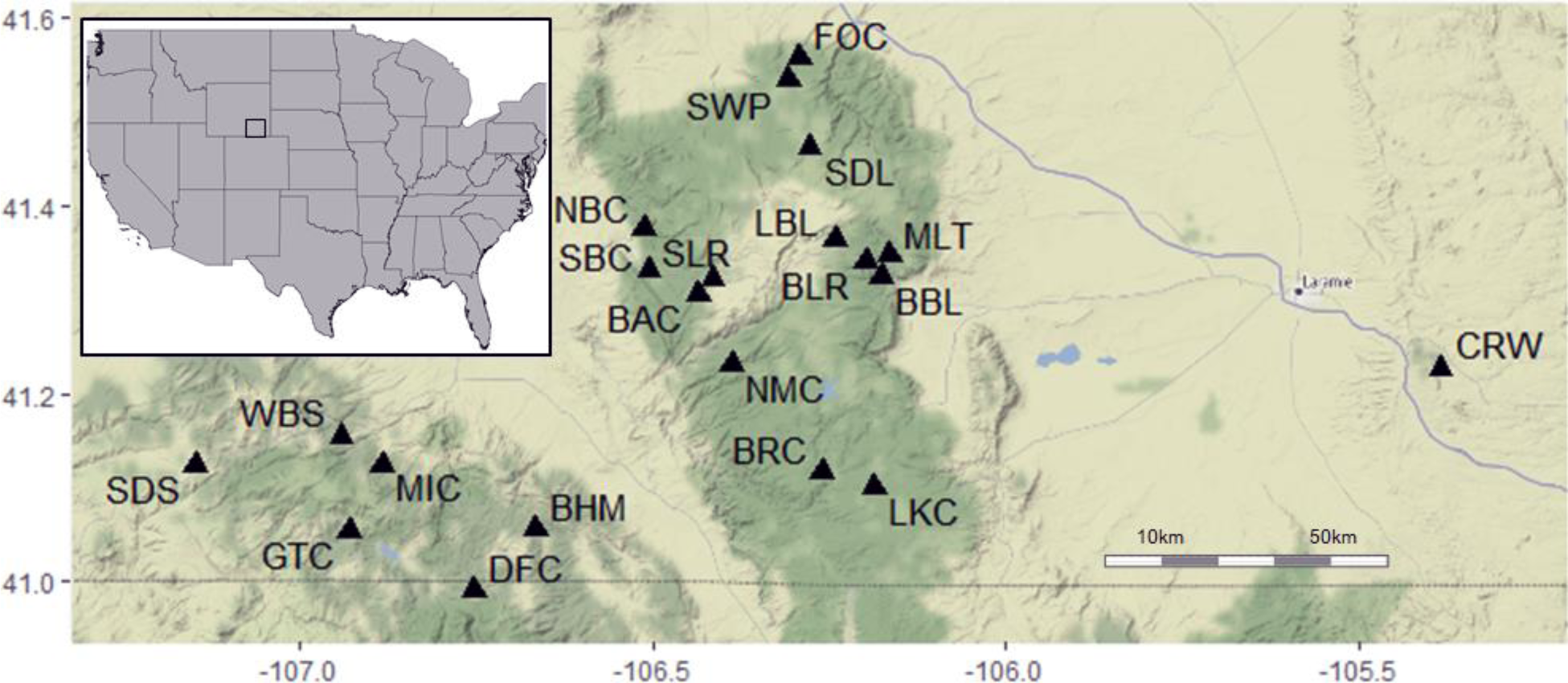
The locations of the 20 populations included in the common garden, along with the Barber Lake (BBL) population which was the source of the 20 maternal family lines, are distributed through the Medicine Bow National Forest. The populations are found in southeastern Wyoming near the University of Wyoming in Laramie, Wyoming, USA.

Several measures of life history transitions, or phenology, were collected on plants in the field. Because *B. stricta* requires vernalization for flowering to occur, the first measures of phenology and size were taken in the summer of 2016, after the plants had over-wintered. Measurements were taken over the entire growing season (April to September) in 2016 through 2018, providing three years of data. We recorded five key phenological timepoints for the individual replicates: time of bolting (defined as the point when the apical meristem differentiated from leaf production to production of a bolting inflorescence and was ∼1cm above the plant rosette); time of flowering (defined as the timing of anthesis of the first flower); time of first fruit initiation (defined as the time when the first silique had formed and was ∼1cm in length); time of first mature fruit (defined as when the first silique was fully formed and the seeds were evident inside the fruit); and time of completion of reproduction (defined as the time when all flowers had completed reproduction and all fruits had matured to produce seeds). We also measured several estimates of size: rosette diameter (defined as the distance between the tips of two leaves on opposite sides of the rosette), longest leaf (defined as the length of the longest leaf measured from the rosette center to the tip of the leaf), and number of rosette leaves at flowering (count of leaves in the basal rosette at flowering). Plant height was determined at first anthesis, at formation of first fruit, and at completion of flowering. The total number of fruits produced was counted at completion of reproduction. After measurements were taken for completion of reproduction, the fruits were collected to harvest the seeds and the inflorescence was removed to simulate natural field condition, in which the inflorescence is typically broken in the winter due to wind/snow.

We used accumulated growing degree days (GDD) to estimate the elapsed time until a given phenological stage was reached. In comparison to calendar days, growing degree days better represent the accumulation of warmth that contributes to growth and developmental transitions in the plants. To determine growing degree days, we used micrometeorological data from the Weather Underground (wunderground.com). Days with mean temperature values above freezing were added each day (in degrees Celsius) to estimate GDD.

Original estimates of population mean circadian period were derived from a larger dataset of 30 populations of *B. stricta* (McMinn *et al*., 2022). Only maternal family lines for each population that were included in the common garden were used to estimate circadian period, thus producing slightly different but strongly correlated (r = 0.80, p < 0.001) mean values per population in comparison to those found in McMinn et al. 2022 (Supplemental Table 1). From the BBL population, 17 of the 20 maternal family lines planted in the common garden had circadian period estimates.

### 2.3 Statistical analysis

We aimed to characterize phenological stages in multiple ways. We estimated the elapsed time to a given stage (e.g., GDD to flowering) for maternal family lines and populations, the extent to which time to a given stage varied among growing seasons (e.g., the degree to which genotypes that flowered early in the first season also flowered early in a subsequent season or showed genotype x environment interactions), and if different stages (e.g., GDD to flowering and GDD to completion of reproduction) were inter-correlated.

For the first set of analyses, we reduced the dataset to replicates that flowered in two of the three years and tested for assumptions of normality and homogeneity of variances for the data using base R (R 4.1.2; R Core Team, 2021). Outliers within a maternal family or population in each year of measurement were determined and removed by the “EnvStats” package (v2.4.0; Millard, 2013). We used two-way mixed ANOVA to partition variation in phenological traits among population, year, the spatial planting block, and the interaction of population × year, with population and year both as fixed factors. Two-way ANOVAs were conducted using the R package “rstatix” (v0.7.0; Kassambara, 2021). The same methods were used to partition variation in phenotypic traits among the maternal families within BBL, testing for variation among maternal family line, year, the spatial planting block, and the interaction of maternal family line × year. Population and maternal family mean values were estimated from the preceding models, and we tested the association between mean population values in phenological traits with elevation at the home site. To understand the contribution of population or family to variation in phenology and circadian period, we estimated variance components using the restricted maximum likelihood method in the lme4 package in R (v1.1-27.1; Bates *et al*., 2015), with population or family as a random factor.

Given an observed elevational cline in phenology (see Results), we were interested in quantifying the association of environmental variables at the home sites and of genotypic circadian period with the expression of life history traits in the common garden. We first assessed spatial autocorrelation and the potential that population values were not independent. Spatial autocorrelation was determined among the population home sites using Moran’s I and Geary’s C to test for spatial clustering of phenology and size in R 4.1.2 packages “sp” (v1.4-5; Bivand *et al*., 2013), “spdep” (v1.1-12; Bivand *et al*., 2013), “rgdal” (v1.5-27; Bivand *et al*., 2021), “spgwr” (v0.6-34 ; Bivand and Yu, 2020), and “spatstat” (v2.2-0; Baddeley and Turner, 2005). Spatial autocorrelation was non-significant, and we therefore proceeded with populations as independent data points. Following a False Discovery Rate control of multiple tests, none of the phenotypic traits tested among the 20 populations showed significant spatial patterning (non-significant for both Moran’s I and Geary’s C), and the populations thus do not appear to be structured by spatial proximity. We therefore treated populations as independent replicates and proceeded with an analysis of environmental variables and their relationship to phenology.

For estimates of climate at the 20 population sites, we utilized data in the WorldClim dataset for 19 bioclimatic variables through R 4.1.2 in the package “raster” (v3.5-2; Fick and Hijmans, 2017; Hijmans, 2021). For the analysis of environmental data, values for the climate variables were collected at 30-s resolution (∼1 km^2^). Soil variables were measured previously at the field sites as was circadian period for each population and maternal family (McMinn *et al*., 2022). We generated principal components for the 27 environmental variables as well as circadian period using the R package “factoextra” (v1.0.7; Kassambara and Mundt, 2020) to reduce the dimensionality, as climatic variables were highly correlated and prior work has shown that circadian period and climate are correlated in *B. stricta* (McMinn *et al*., 2022).

Bivariate correlations estimated between individual plant phenotypes (e.g., life history traits) using the R package “corrplot” (v0.91; Wei and Simko, 2021) were strongly positive. Variance inflation factor in the “car” package (v3.0-12; Fox and Weisberg, 2019) tests revealed significant multicollinearity among phenology traits. Therefore, we reduced dimensionality of phenotypic data through principal component analysis (package “factoextra” v1.0.7) for each of the three years of measurement.

We used multivariate linear regression of the phenotypic PCs on the environmental PCs to assess the extent to which plant phenotypes were associated with climate or soil conditions in the site of origin or with population values of circadian period. We further evaluated associations specifically between plant phenotypes and environmental features using principal component regression (PCR) and partial least squares regression (PLS). PCR first computes the principal components of the predictors, and then uses these components as predictors in a regression against the response variable (Jolliffe 1982). PLS regression is a similar analysis to PCR but works in a supervised framework, as the components are informed by the dependent variables and dimensionality is reduced with a focus on covariance (Wold *et al*., 2001). Variables were standardized by dividing each by their standard deviation. The model with the best explanatory power was evaluated via cross-validation both within each model and as a comparison of the PLS and PCR models. After the optimal model was determined, we calculated the contribution of each coefficient. For this analysis, we used R 4.1.2 and packages “pls” (v2.8-0; Mevik *et al*., 2020) and “caret” (v6.0-9.0; Kuhn, 2020).

Because circadian period was not heavily weighted on any of the predictor PCs, we also performed regressions of individual phenological traits on circadian period. We subsequently estimated multi-year mean values for phenological points (GDD to bolting, to flowering, to first fruit, to mature fruit, to reproductive completion) as the geometric mean for each trait over all three years (e.g., geometric mean for GDD to flowering in yr 1, 2, and 3), and regressed these geometric estimates on circadian period to test consistency of phenology-clock associations among years in the face of inter-year environmental variability.

Fitness estimates were determined both for the plant populations and the maternal families within each year. We used fecundity (estimated as total fruit number) as one component of fitness along with overall survival and then utilized two analyses to examine potential drivers for plant fitness. First, we wanted to assess the effect of environmental divergence between the common garden and the 20 homesites of the populations. We determined mean annual temperature and precipitation by elevation (WorldClim data) and estimated the difference between conditions in each of the 20 population sites vs. the common garden location. We used linear regression with the environmental differences as our predictor of geometric mean lifetime fruit production and overall survival for each population, under the hypothesis that greater environmental differences between the home site vs. the experimental common garden site would reduce fitness. Second, to better understand which traits were directly or indirectly affecting fitness, we utilized structural equation modeling. Because fewer plants flowered in the first year, we used the second and third years’ data in the model. Starting with a fully saturated model with all developmentally possible relationships, we used year 3 data to both identify and remove traits with non-significant explanatory power, i.e., identify a best-fit model. The reduced model was evaluated with the year 2 measurements. We hypothesized that circadian period would influence both phenology and plant size and that these plant traits would directly affect the number of fruits produced (Figure 7), i.e., circadian period would exert indirect selection on plant fitness. Based on prior studies in *B. stricta* (Anderson *et al*., 2011), the length of vernalization is an important cue for flowering; hence we further hypothesized that the timing of flowering could affect size measures (height at flowering, rosette and cauline leaf number). Plant height (at flowering and the final measured height), the number of rosette leaves, and the number of cauline leaves were included as correlated factors. Trait values were standardized in the model, and traits with significant explanatory power are presented with the overall statistical estimates of the model fit. For these analyses, we used R 4.1.2 with the packages “lavaan” (v.0.6-11; Rosseel, 2012), “semPlot” (v1.1.4; Epskamp, 2022), and “DiagrammeR” (v1.0.9; Iannone, 2022).

## Results

### 3.1 Variation in traits among and within populations

Many phenological traits demonstrated significant differences by population, year, and the interaction of population ⨯ year (Table 1, Figure 2). Growing degree days to all phenological time points (GDD to bolting, flowering, first fruit initiation, first mature fruit, and completion of reproduction) was significantly different among the 20 populations; only 1 of 4 intervals between developmental points (Interval of bolting to flowering) showed significant differences among populations. Year, or growing season, significantly affected time to all phenological timepoints and all phenological intervals, with the effect of year differing between early *vs*. later developmental stages. In particular, GDD to phenological timepoints expressed early in the life history (to bolting, to flowering, and to first fruit, Figure 2a-c) was delayed in year 1 relative to year 2, but this delay was absent for later phenological traits (GDD to mature fruit, Figure 2d) and reversed rank order for GDD to complete reproduction (Figure 2e), indicating that while early stages were delayed under environmental conditions in year 1 later stages were not (Figure 2). More specifically, the difference in GDD to bolting was 200 degree days between years 1 and 2, while for GDD to flowering it was only 150, for first fruit it was 75, and for GDD to mature fruit there was no difference between years 1 and year 2. For GDD to complete reproduction, year 2 was delayed relative to year 1 by ∼75 degree days. GDD to all phenological stages showed a delay in year 2 relative to year 3. The total number of fruits at the end of the season was higher in years 2 and 3 than year 1. The interaction of population × year significantly affected most phenologic timepoints (GDD to flowering, first fruit production, and completion of reproduction) as well as intervals between some timepoints (interval of bolting to flowering and mature fruit to completion of reproduction; Table 1, Supplemental Figure S1). All measurements of size varied significantly by population, year, and their interaction (Table 1). The significant main effect of population indicates that a trait’s phenotypic expression in one year was positively correlated with the phenotype expressed in another growing season, while the significant population x year interaction indicates that the across-season correlation was < 1. Correspondingly, phenological traits were significantly positively correlated within and across years (r = 0.54 to 0.98 for all pairwise correlations).

**Figure 2.**
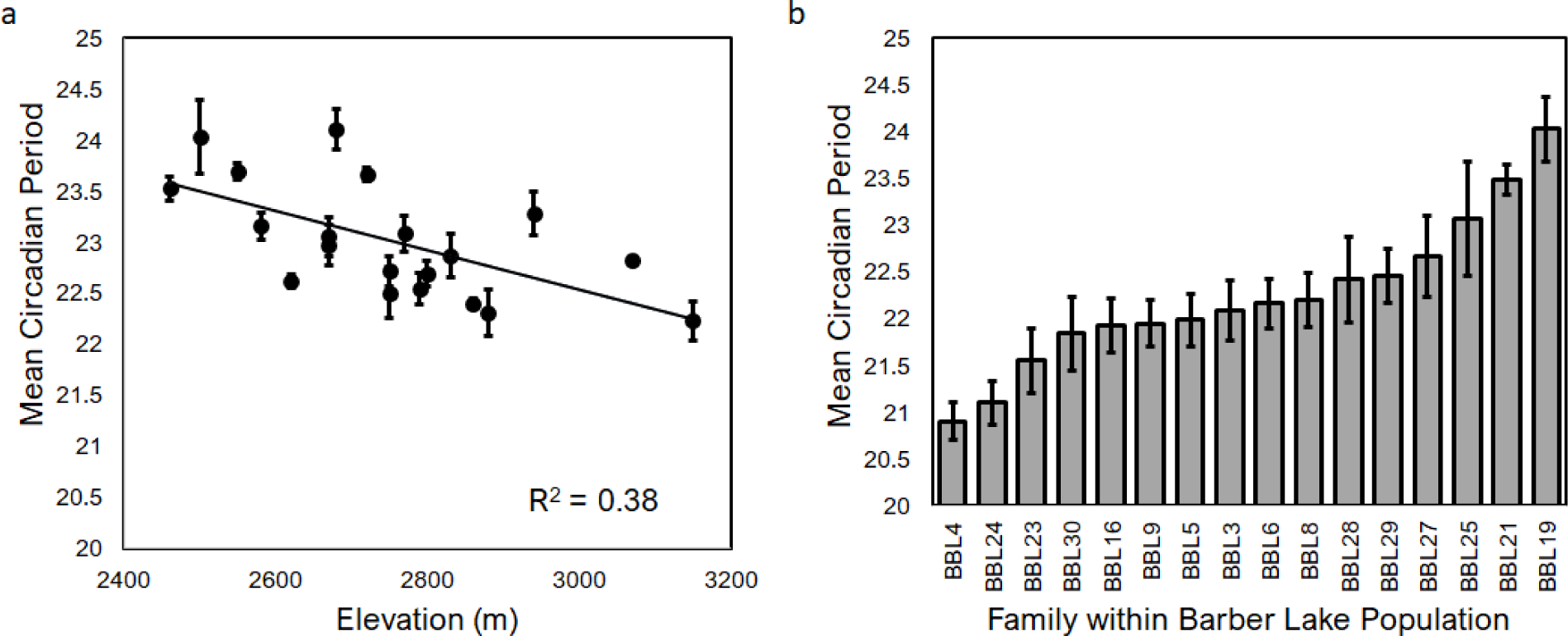
Variation in populations for growing degree days to different phenological points across the three years of measurement. Significant differences among years for the traits is shown by letters at the top of each panel (a, b, or c). Grey lines indicate population mean value, and the black line indicates mean value over all populations.

**Table 1.**
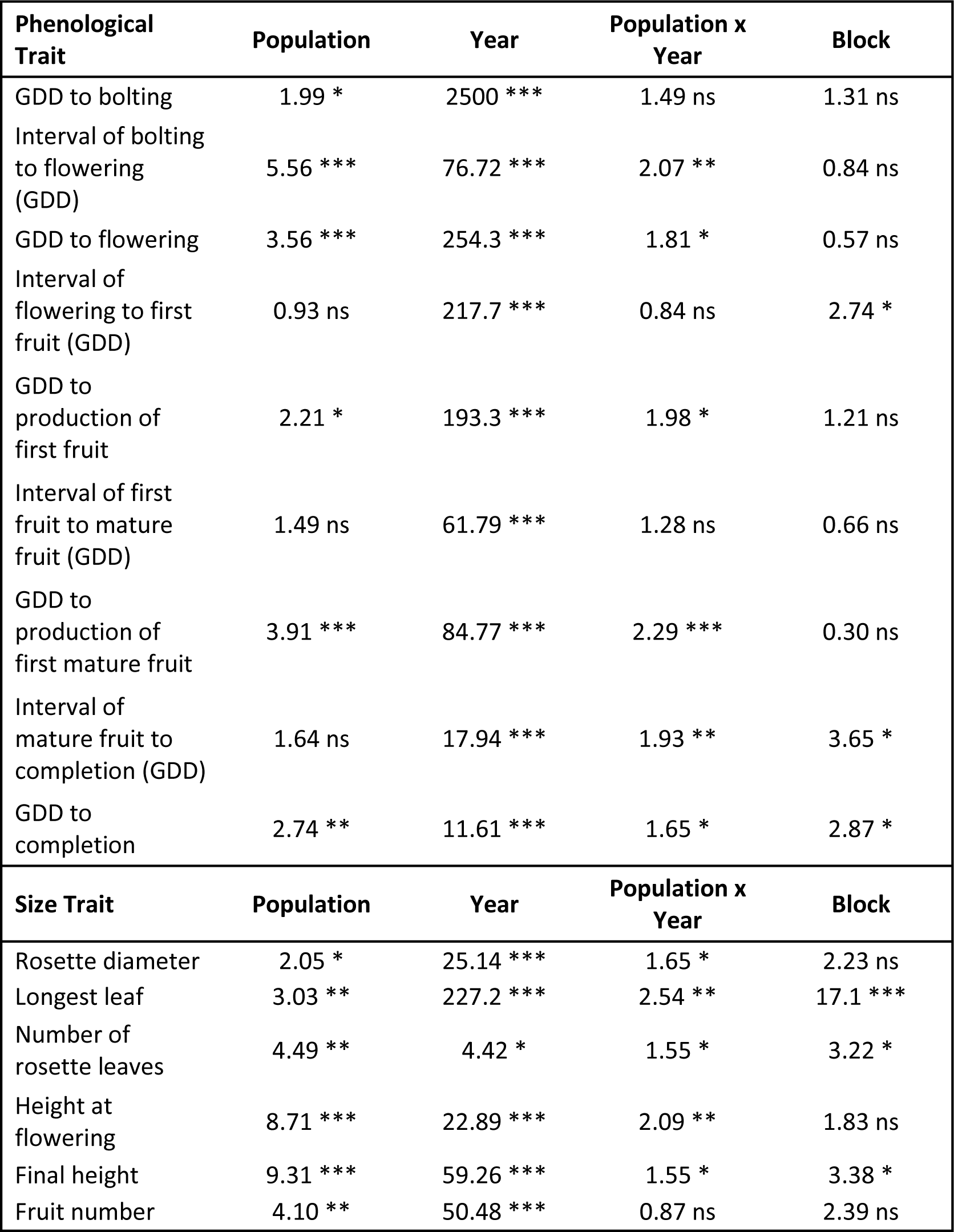
ANOVA testing the effects of population, year, and the interaction of population and year on phenology and plant size in the dataset with 20 populations. Values in the table reflect F-test statistic and associated p-values.

| **Phenological Trait** | **Population** | **Year** | **Population x Year** | **Block** |
| --- | --- | --- | --- | --- |
| GDD to bolting | 1.99 * | 2500 *** | 1.49 ns | 1.31 ns |
| Interval of bolting to flowering (GDD) | 5.56 *** | 76.72 *** | 2.07 ** | 0.84 ns |
| GDD to flowering | 3.56 *** | 254.3 *** | 1.81 * | 0.57 ns |
| Interval of flowering to first fruit (GDD) | 0.93 ns | 217.7 *** | 0.84 ns | 2.74 * |
| GDD to production of first fruit | 2.21 * | 193.3 *** | 1.98 * | 1.21 ns |
| Interval of first fruit to mature fruit (GDD) | 1.49 ns | 61.79 *** | 1.28 ns | 0.66 ns |
| GDD to production of first mature fruit | 3.91 *** | 84.77 *** | 2.29 *** | 0.30 ns |
| Interval of mature fruit to completion (GDD) | 1.64 ns | 17.94 *** | 1.93 ** | 3.65 * |
| GDD to completion | 2.74 ** | 11.61 *** | 1.65 * | 2.87 * |
| **Size Trait** | **Population** | **Year** | **Population x Year** | **Block** |
| Rosette diameter | 2.05 * | 25.14 *** | 1.65 * | 2.23 ns |
| Longest leaf | 3.03 ** | 227.2 *** | 2.54 ** | 17.1 *** |
| Number of rosette leaves | 4.49 ** | 4.42 * | 1.55 * | 3.22 * |
| Height at flowering | 8.71 *** | 22.89 *** | 2.09 ** | 1.83 ns |
| Final height | 9.31 *** | 59.26 *** | 1.55 * | 3.38 * |
| Fruit number | 4.10 ** | 50.48 *** | 0.87 ns | 2.39 ns |

For maternal family lines within the BBL population, variation for phenological traits approximately resembled patterns observed among the 20 populations (Table 2, Supplemental Figure S2). Maternal family lines differed significantly for four of the five phenological timepoints (all but GDD to bolting). As for populations, the year of measurement significantly affected the GDD to all phenological stages. Earlier phenological stages (GDD to bolting, to flowering, and to first fruit) again displayed a greater delay in timing than did later phenological stages (GDD to mature fruit and completion of reproduction) in year 1 relative to year 2 (Supplemental Figure S2). Maternal family × year interactions were of lesser magnitude and less frequent than were main effects of maternal family and year (Table 2). Intervals between stages in the maternal family lines were delayed in later years but showed the greatest length between fruiting and completion stages in the second year (Supplemental Figure S3). Both maternal family and year significantly affected estimates of plant size (Table 2).

**Table 2.** ANOVA for families of the Barber Lake population on plant phenology and size. Values reflect F-test statistic and associated p-values.

| **Phenological Trait** | **Maternal Family** | **Year** | **Maternal Family x Year** | **Block** |
| --- | --- | --- | --- | --- |
| GDD to bolting | 1.28 ns | 843.5 *** | 0.96 ns | 3.69 * |
| Interval of bolting to flowering (GDD) | 2.77 ** | 44.22 *** | 1.27 ns | 1.14 ns |
| GDD to flowering | 3.05 ** | 1143.3 *** | 2.29 * | 0.81 ns |
| Interval of flowering to first fruit (GDD) | 0.99 ns | 47.98 *** | 1.19 ns | 1.07 ns |
| GDD to production of first fruit | 2.08 ** | 100.1 *** | 1.98 * | 1.15 ns |
| Interval of first fruit to mature fruit (GDD) | 3.76 * | 117.7 *** | 0.80 ns | 3.47 ** |
| GDD to production of first mature fruit | 4.24 ** | 32.86 *** | 2.12 * | 5.83 ** |
| Interval of mature fruit to completion (GDD) | 0.78 ns | 19.05 ** | 1.02 ns | 1.29 ns |
| GDD to completion | 3.53 ** | 18.42 *** | 1.97 * | 5.04 ** |
| **Size Trait** | **Maternal Family** | **Year** | **Maternal Family x Year** | **Block** |
| Rosette diameter | 4.37 ** | 29.72 *** | 1.20 ns | 8.07 ** |
| Longest leaf | 2.41 ** | 162.2 *** | 1.65 * | 7.72 ** |
| Number of rosette leaves | 4.97 ** | 40.29 *** | 1.43 ns | 5.29 * |
| Height at flowering | 2.80 ** | 22.87 *** | 1.59 * | 4.47 * |
| Final height | 3.52 ** | 52.98 *** | 1.29 ns | 4.26 * |
| Fruit number | 1.97 * | 35.46 ** | 0.87 ns | 3.68 * |

Year 2 had the largest proportion of flowering individuals (83% of surviving plants), and we therefore looked within that year to compare the relative magnitude of variation segregating among experimental populations vs. among families within the BBL population. The variance component for the effect of maternal family for the trait of GDD to flowering explained 8% of the total variation, while the variance component for the population effect explained 18% of the total variation. This indicates that overall, a greater proportion of the variation for phenology was found among populations than within the single population. Two populations expressed exceptionally high intrapopulation ranges of GDD to flowering (Green Timber Creek, GTC, and Sandstone, SDS; Supplemental Table S1); both populations are located in the western mountain range of the Sierra Madres (Supplemental Table S1).

Mean circadian period differed significantly among populations used here (F_19_=18.07, p<0.001) and displayed a range of ∼2h, from 22.23h to 24.11h (Figure 3a; Supplemental Table S1). The populations used here showed an elevational cline, in which circadian period duration was longer for populations from low elevation and shorter for high elevation populations. Maternal families within the BBL population also differed significantly in circadian period length (F_15_=3.15, p<0.001), showing a 3h range from 20.91h to 24.02h (Figure 3b; Supplemental Table S2).

**Figure 3.**
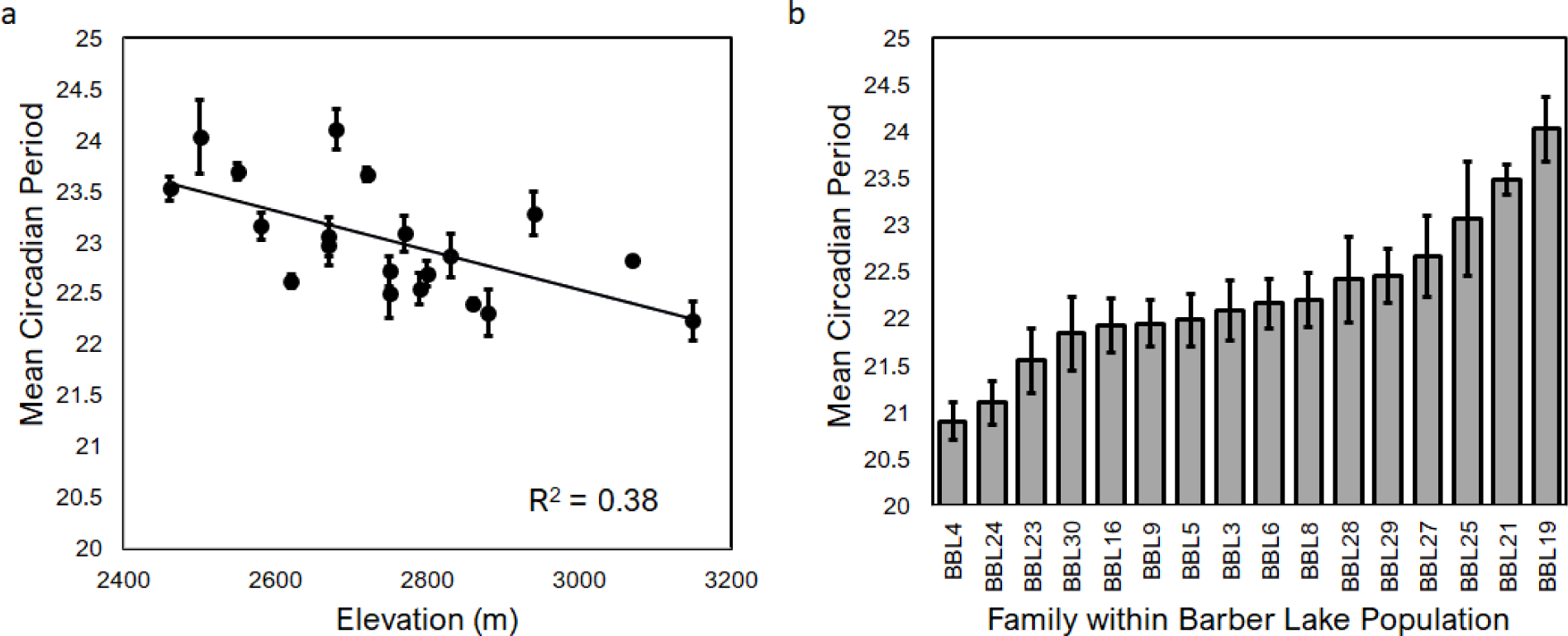
Mean circadian period for populations (a) and maternal families within the Barber Lake population (b). Populations are graphed against the elevation of the home site. Standard error bars are included.

### 3.2 Environmental association with measured traits

All phenological traits (e.g. GDD to flowering) exhibited a significant negative correlation with elevation (Figure 4a). To understand what environmental variables could be contributing to this elevational cline in phenology, we reduced multivariate environmental and inter-correlated phenotypic data and tested for their association. First, we reduced data of predictor variables, including climate data and circadian period data, via Principal Components Analysis (and refer to these as “environmental PCs”). The first five environmental components explained ∼93% of the variation. The first environmental PC (43.8% of the variation) was heavily weighted by mean annual temperature, temperature during the growing season, and annual precipitation (Supplemental Table S3). The second environmental PC (22.3% of the variation) was heavily weighted by metrics of interannual temperature and precipitation variability. The circadian period was most heavily weighted on PC1 (although this weighting of 0.55 was lower than the top variables).

**Figure 4.**
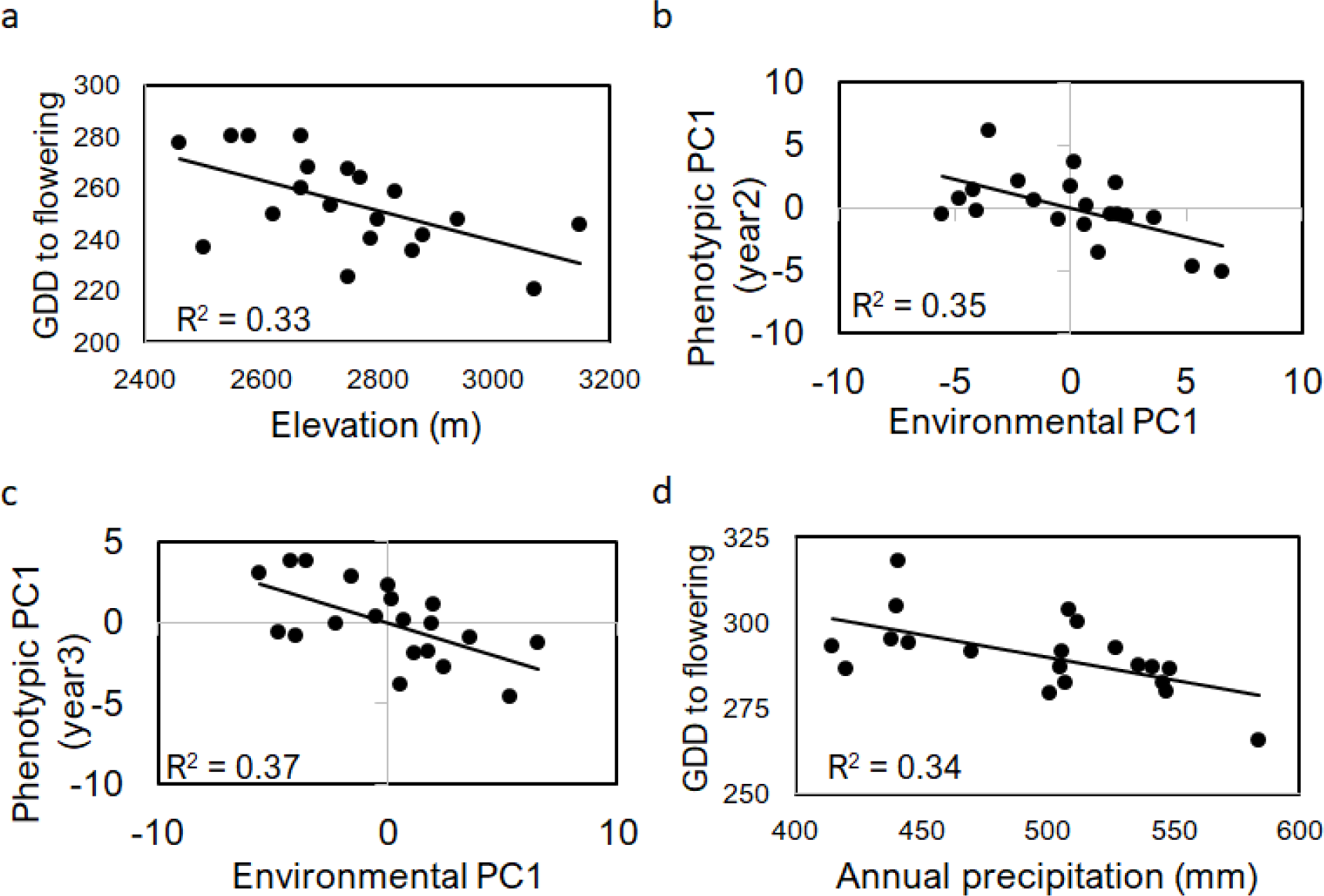
Elevation as a single variable explains phenological differences for the populations in the common garden (panel a). Based on PLS regression, annual precipitation is the strongest environmental predictor for phenological measurements of the populations of *Boechera stricta*. The phenological values for populations in Year 2 are shown as a univariate response to annual precipitation and the negative relationship (time to phenological points increase with reduced precipitation) is presented in subsequent stages.

We reduced the plant trait data (phenology and size) with PCA for each of the three years of measurement (and refer to these as “phenotypic PCs”) (Supplemental Table S4). For all three years, the first phenotypic PC was heavily weighted by GDD to flowering, to first fruit, to mature fruit, and to complete reproduction, whereas the second phenotypic PC was weighted by the height at flowering and final height and the diameter of the plant rosette at flowering (Supplemental Table S4). The first two environmental PCs were used as predictor variables for the first two phenotypic PCs in multivariate linear regression. In the first year, the overall model was not significant for either of the phenotypic PCs. In both the second year and third years, the environmental PCs were significantly associated with phenotypic PC1 (year 2: F=14.29; year 3: F=7.74; Table 3). Environmental PC1 showed a negative relationship with phenotypic PC1 in year 2 (Figure 4b) and year 3 (Figure 4c), indicating that lower annual and growing season temperatures, as well as higher annual precipitation and shorter circadian period led to earlier plant phenology. Environmental PC2 showed a moderately significant negative association with phenotypic PC1 in year 2 only (Table 3), indicating that increased diel and interannual variability in temperature and reduced seasonality of precipitation were associated with delayed plant phenology.

**Table 3.**
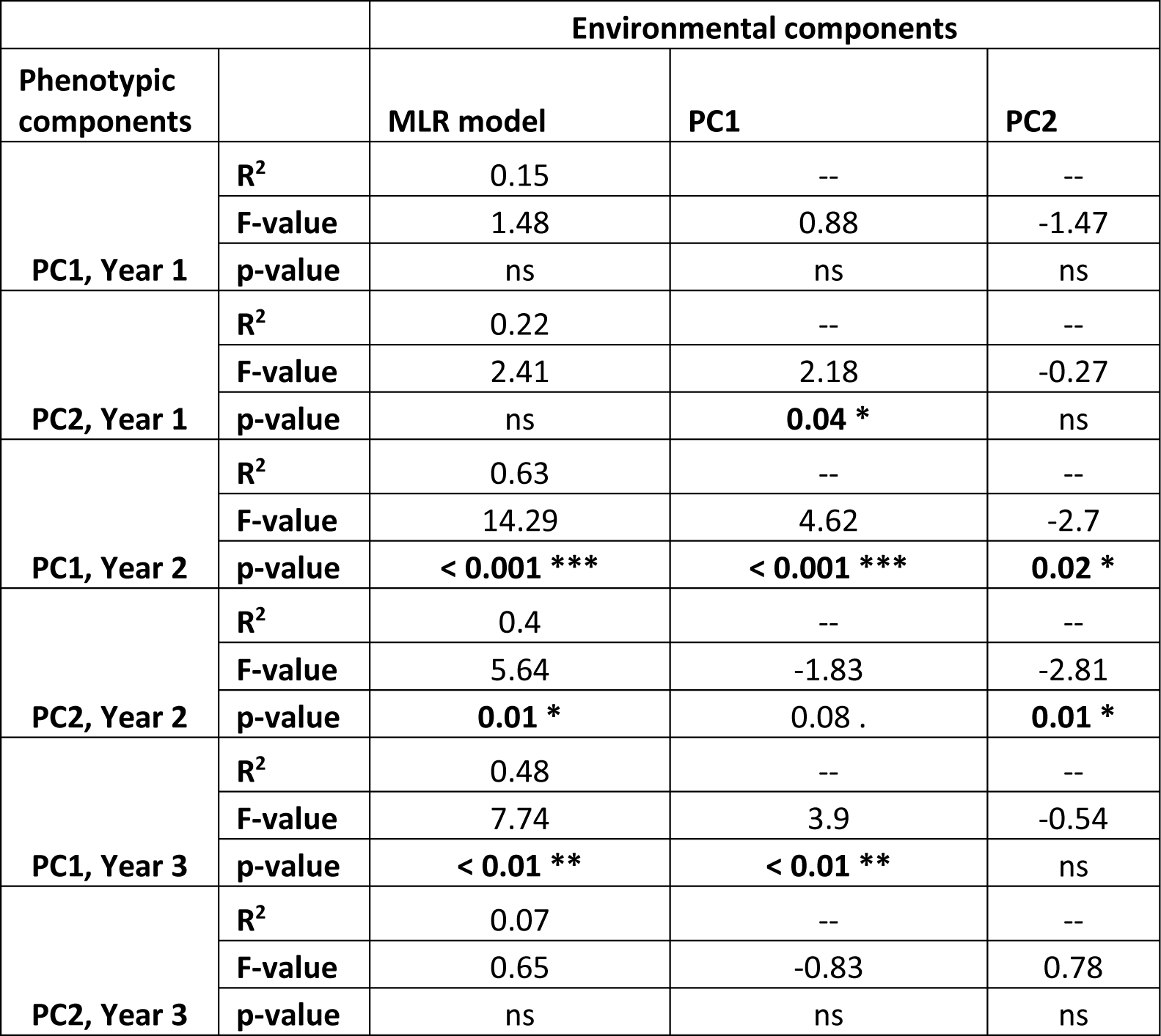
Multiple linear regression results show the combined effect of the first two components of the environmental PCA on the components of the plant traits. The full model R^2^ and the F-test statistics for the full model and each individual component are shown with the level of significance.

|  |  | Environmental components |  |  |
| --- | --- | --- | --- | --- |
| Phenotypic components |  | MLR model | PC1 | PC2 |
| PC1, Year 1 | $R^2$ | 0.15 | -- | -- |
|  | F-value | 1.48 | 0.88 | -1.47 |
|  | p-value | ns | ns | ns |
| PC2, Year 1 | $R^2$ | 0.22 | -- | -- |
|  | F-value | 2.41 | 2.18 | -0.27 |
|  | p-value | ns | **0.04 \*** | ns |
| PC1, Year 2 | $R^2$ | 0.63 | -- | -- |
|  | F-value | 14.29 | 4.62 | -2.7 |
|  | p-value | **< 0.001 \*\*\*** | **< 0.001 \*\*\*** | **0.02 \*** |
| PC2, Year 2 | $R^2$ | 0.4 | -- | -- |
|  | F-value | 5.64 | -1.83 | -2.81 |
|  | p-value | **0.01 \*** | 0.08 . | **0.01 \*** |
| PC1, Year 3 | $R^2$ | 0.48 | -- | -- |
|  | F-value | 7.74 | 3.9 | -0.54 |
|  | p-value | **< 0.01 \*\*** | **< 0.01 \*\*** | ns |
| PC2, Year 3 | $R^2$ | 0.07 | -- | -- |
|  | F-value | 0.65 | -0.83 | 0.78 |
|  | p-value | ns | ns | ns |

To further define which environmental factors most strongly contribute to the environment PC1-phenotype PC1 association observed in years 2 and 3, we evaluated PLS and PCR models of the 5 phenological traits with the strongest effect of population (GDD to bolting, flowering, first fruit, mature fruit, completion of reproduction) for each of the two years. Models were compared to determine the strongest predictor variables for the response. For each phenological timepoint, the best fit PLS model included a single predictor component, accounting for 94.45 – 99.96% predictor variation and 26.94 – 71.30% of the variation in the response variables. Across the 10 PLS and PCR models (5 phenological response variables × 2 years), many of the same variables (mean annual temperature, temperature during the growing season, and annual precipitation) from the PCA models above were strongly weighted in the best-fit models. The PLS and PC regression compared the models with the most explanatory power and determined that annual precipitation was the strongest predictor of late phenology for both years 2 and 3, and in particular higher annual precipitation at the home site of each population was associated with earlier phenology (Figure 4d). In sum, both univariate linear regression with principal components of the environment variables as well as PLS/PC regression models indicate the negative relationship of phenology and annual precipitation.

### 3.3 Association of circadian period with phenology

To characterize the potential relationship between phenological trait expression and circadian clock function, we examined the relationship between individual phenological traits and circadian period. Because the environmental PC1 (i.e., the PC weighted with circadian period) was only associated with phenotypic PC1 in years 2 and 3, we focused on those years of data. In the second year of measurements for the populations, circadian period was positively associated with phenotypic PC1 (F=8.02, r=0.52, p=0.01; Figure 5a), which was weighted by GDD to flowering (Figure 5b), to first fruit, to mature fruit, and completion of reproduction (data not shown); GDD to bolting was notably unrelated to circadian period (Figure 5c). Maternal families showed a similar pattern of phenological associations with circadian period in the year 2 measurements, again with a notably strong association between GDD to flowering and circadian period (Supplemental Figure S4). Circadian clock-phenological associations were the same in year 3 (data not shown).

**Figure 5.**
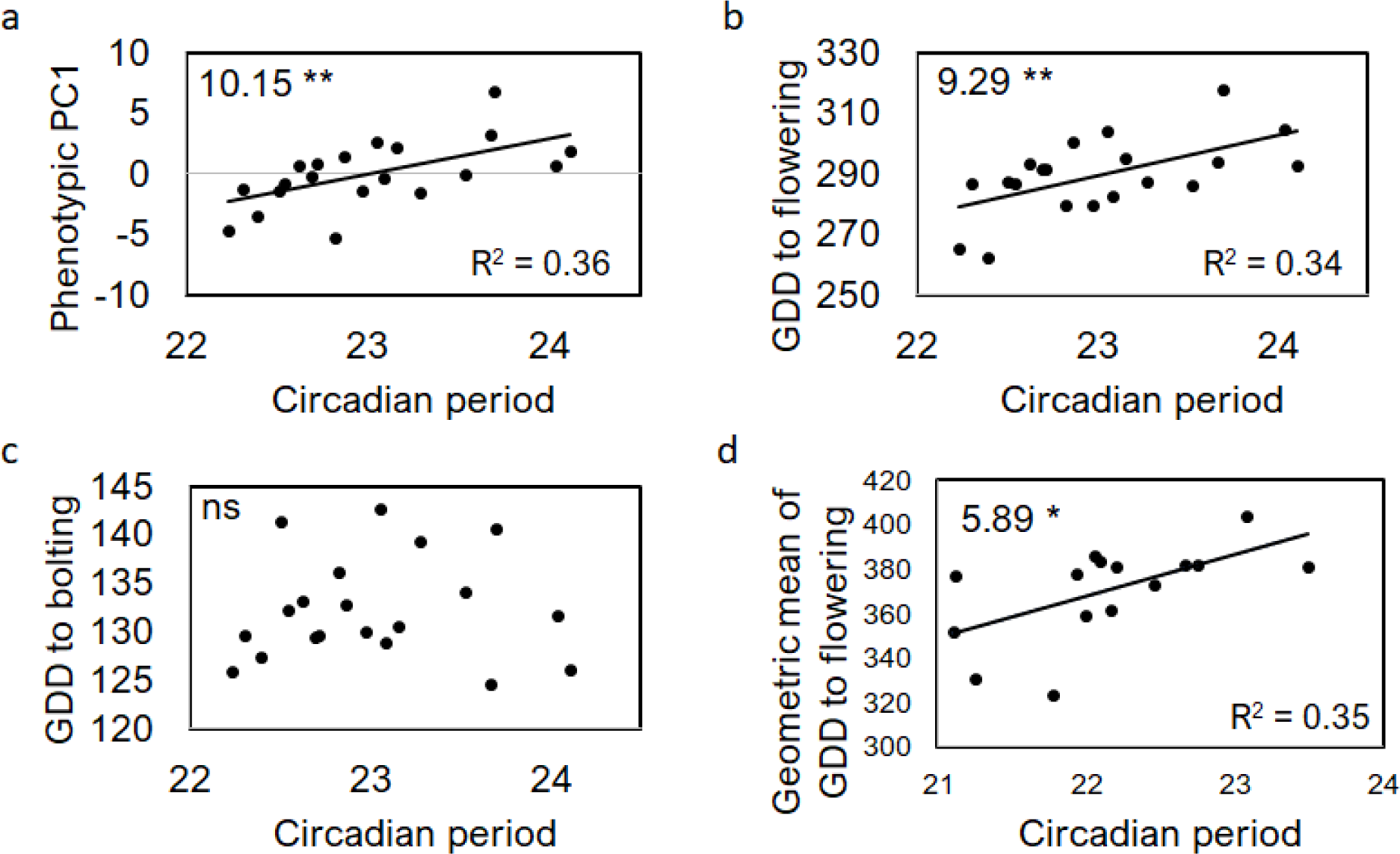
Measurements for phenology from the populations as explained by circadian period. In the second year of measurements, phenotypic PC1 (weighted by phenology) is strongly associated with circadian period (panel a). Growing degree days to flowering in the second year shows an association with circadian period (panel b), but the time to bolting does not (panel c). The geometric mean across all 3 years for GDD to flowering also correlates with circadian period (panel d). F-values, r-squared values, and level of significance are included on the panels.

We used the geometric mean over all three years of individual phenological traits to assess consistency of associations with period in the face of inter-annual variability. Among populations, circadian period was found to have a positive association with the geometric mean of GDD to flowering (Figure 5d), first fruit, mature fruit, and completion of reproduction among the populations (Supplemental Figure S5). The positive relationship between the geometric mean of phenology and circadian period was also evident in the maternal family lines (all significant relationships for GDD to flowering, first fruit, mature fruit, and complete reproduction; r=0.43 to 0.58, Supplemental Figure S6). With a longer circadian period, the time required for plants to flower and produce fruits is delayed.

### 3.4 Indirect selection for circadian period

Testing the fitness consequences of environmental differences between the home site of each population and the common garden site, we found that elevation and mean annual temperature did not have an effect. However, mean annual precipitation had a significant negative effect (Figure 6); plants from wetter environments had reduced lifetime fruit production and reduced survival at the drier common garden location.

**Figure 6.**
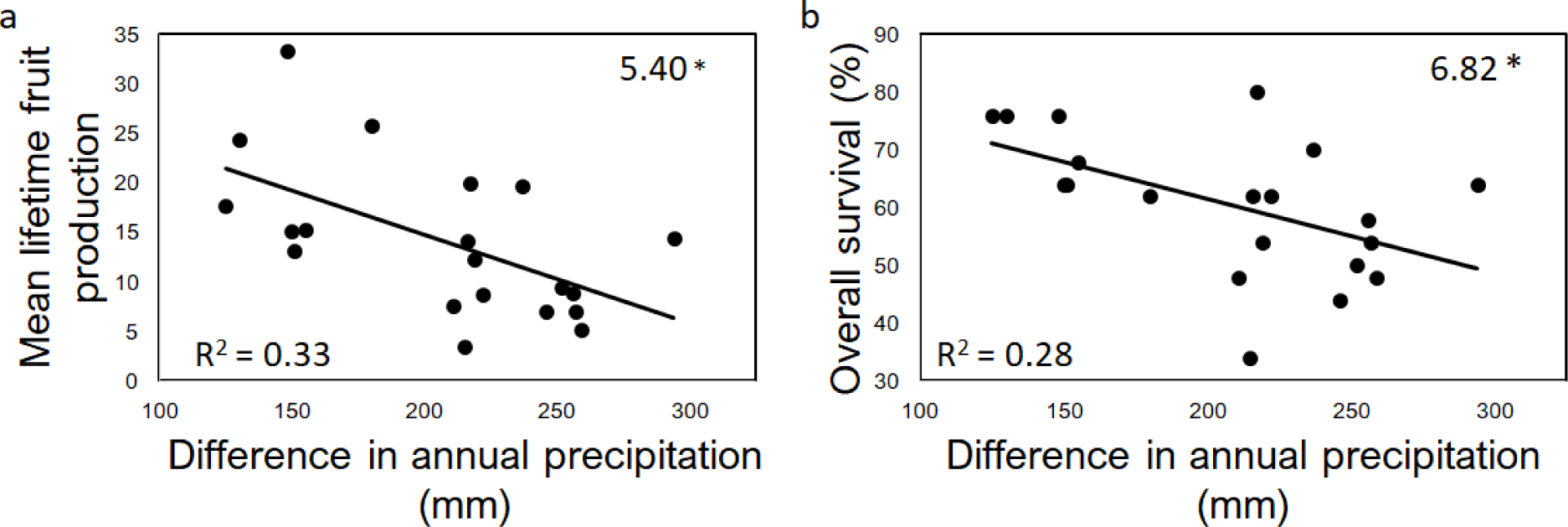
Increased difference in the amount of annual precipitation at the site of origin for the 20 populations from the annual precipitation in the common garden locations shows a negative correlation with fecundity (panel a) and survival (panel b) in the plants.

We used year 3 data to estimate the best fit structural equation model, that is, the combination of phenotypic traits that best explained variation in the fitness component of fruit number. The estimated model for the dataset indicates a good fit of the data based on model values for chi-square statistic, comparative fit index, and root mean square of approximation (Table 4). The SEM suggests natural selection on size, phenology, and circadian clock timing (Figure 7b). With regard to size (estimated as cauline leaf number, rosette leaf number, height at flowering, and final height), direct selection is positive favoring larger size as evidenced by significant paths from each of the four size traits to fruit set. For measures of phenology (GDD to flowering and GDD to first fruit), direct selection favors early flowering in combination with delayed production of the first fruit, as evidenced by the significant negative and positive partial regression coefficients, respectively, on the path from each trait to fruit set. Indirect selection for the circadian clock through early flowering favors an accelerated circadian period whereas indirect selection through longer time of fruit set favors a decelerated circadian period.

**Table 4.** Model statistics for each of the constrained models from the second year of measurements for path analysis. Full model statistics include the chi-square value with degrees of freedom, root mean square of approximation (RMSEA; minimum, mean, and maximum values), and comparative fit index (CFI). Coefficient values are shown for each variable and its direct effect on fitness as determined by fruit number. In the table, “—” denotes variables that were tested but removed from the constrained model.

|  | **Populations** | **Maternal Families** |
| --- | --- | --- |
| **$\chi^2$ test** | 2566.9, df=17 *** | 1787.2, df=17 *** |
| **RMSEA** | 0.593 ≤ **0.612** ≤ 0.632 *** | 0.541 ≤ **0.563** ≤ 0.585 *** |
| **CFI** | 0.416 | 0.533 |
| **Circadian period** | ns | ns |
| **GDD to flowering** | -0.251 *** | -0.155 * |
| **GDD to fruiting** | 0.101 * | 0.338 *** |
| **GDD to mature fruit** | -- | -- |
| **GDD to complete reproduction** | -- | -- |
| **Rosette leaf number** | 0.217 ** | ns |
| **Cauline leaf number** | 0.199 ** | -0.271 ** |
| **Height at flowering** | 0.203 ** | 0.252 ** |
| **Height at completion** | 0.313 *** | 0.582 *** |

**Figure 7.**
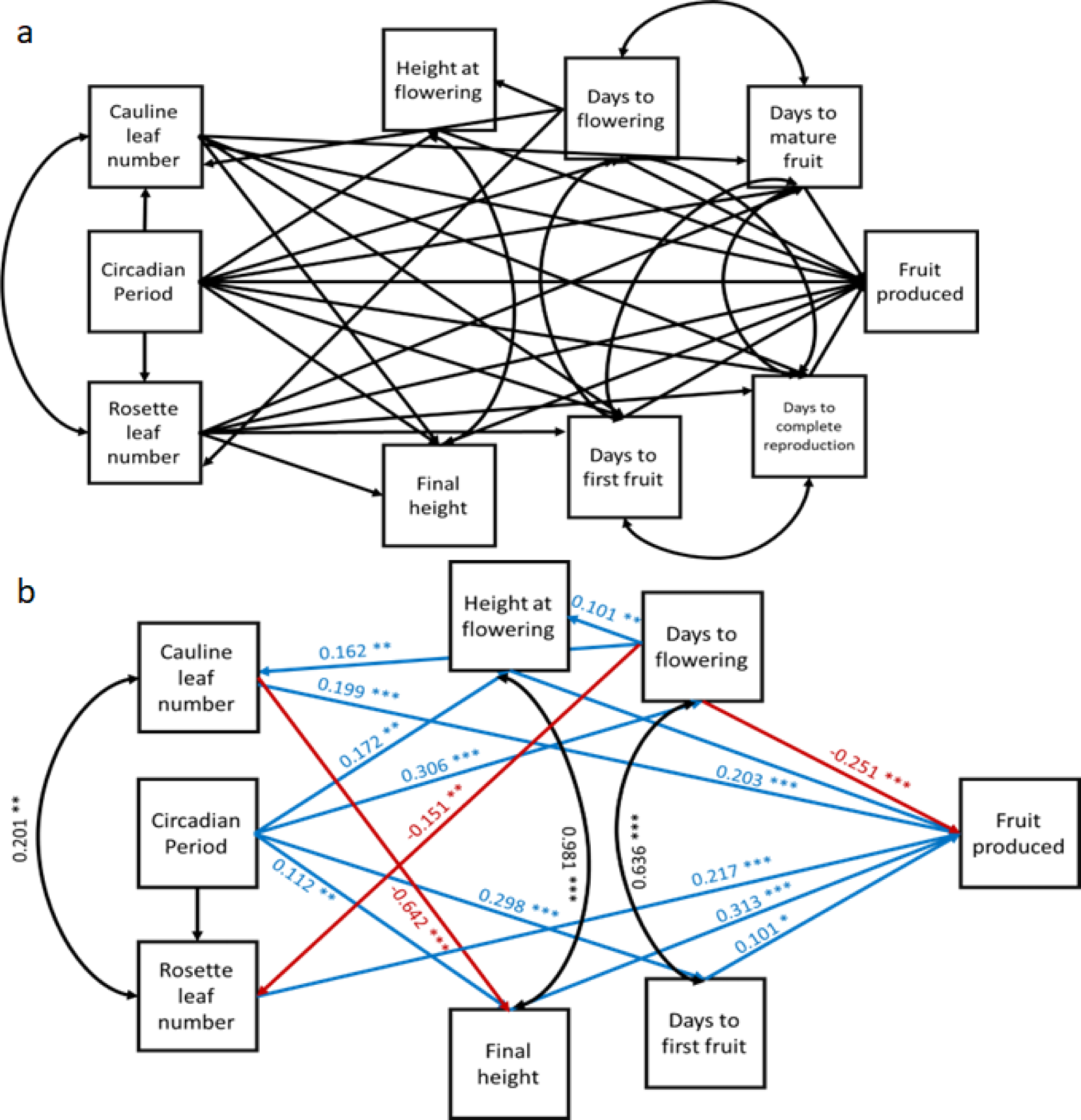
Saturated model of hypothesized relationships contributing to fecundity (panel a). Best-fit model for the second year of population measurements (panel b). Blue lines indicate positive relationships, red lines indicate negative relationships, and curved black lines are covariances. Values for coefficients are included along with significance level for each relationship.

## Discussion

The circadian clock is a critical time-keeping mechanism thought to enable adaptive responses to temporal environmental changes; yet, important aspects of its genetic architecture and multi-year fitness consequences that underlie evolutionary potential remain largely undescribed. Quantitative genetic studies routinely reveal high levels of segregating variation within and among populations (examples in Spitze, 1993, Vitasse *et al*., 2009, Johnston *et al*., 2022). We have previously observed significant segregating variation in circadian clock period both among and within populations of the short-lived perennial, *Boechera stricta*. Here in a multi-year study, we observe significant genetic variances in multiple phenology traits as well as significant environmental associations. Within each experimental year, genetic variance components for phenological traits within the BBL population ranged from 2-11%, and among populations the range was 8-19%, with lower genetic variances in yr 1 and the highest genetic variances in yr 2. Previously, we found population structure for circadian period among 30 local populations of *B. stricta* (McMinn *et al*., 2022). The variance components were larger for circadian period in that study; nevertheless, as for phenology, a greater portion of the variance was partitioned among populations (24%) than within populations (21%) for circadian period. We find that circadian period is an important predictor of plant phenology across multiple growing seasons, and thereby influences fitness; further, variable selection on flowering time across elevations or variable selection on different phenologic stages (flowering time vs. fruiting time) may in part explain the maintenance of genetic variation in circadian clock parameters.

### 4.1 Environmental conditions at the home-site of populations and in the common garden affect phenology

Common-garden studies are frequently used to test for genetic differentiation among populations, potentially indicative of adaptive evolution. Studies in tree species have demonstrated elevational clines in phenological traits among populations (e.g. Salmela *et al*., 2013; Dantec *et al*., 2015; Bucher *et al*., 2020). While the existence of an elevational gradient may be intuitive, it is not a ubiquitous pattern across plant populations. Halbritter *et al*. (2018) reviewed 70 common garden experiments and found a relationship in plant size based on elevation, but the pattern was not present for phenology. Consistent with prior observations in *B. stricta* (Anderson *et al*., 2015), we observed an elevational cline in the expression of phenological traits, such that populations from lower elevations flowered later in the growing season. We found among the *B. stricta* populations that mean annual temperature, annual precipitation, and variation for both temperature and precipitation during the warmest months (the growing season) were strong predictors for phenology. The growing season is longer for *B. stricta* plants coming from lower-elevation sites and allows increased time for flowering and completion of subsequent phenological stages in comparison to higher-elevation sites. In particular, high elevation sites are cooler and wetter with a reduced growing season duration; even when populations were located only a few kilometers apart, elevational differences contributed to significant environmental divergence in local conditions and an elevational cline in phenologic timing was observed.

Beyond genetic differentiation, phenological timing was also sensitive to ambient seasonal conditions in the common garden. We observed a substantial delay in the initiation of reproduction (bolting) in the first year. Climate conditions at the experimental site were not particularly different among growing seasons observed here; for instance, temperatures differed by only 3 degrees across the three years. Rather than environmental effects, the year 1 phenological delay could be due instead to low resource accumulation, that is, low levels of biomass accumulated early in the life history (Van Noordwijk and De Jong, 1986; Roff and Fairbairn, 2007). Interestingly, the phenological delay observed in year 1 for time to bolting was not observed for later phenological stages such as the timing to mature fruit production. Further, for the growing degree days to completion of reproduction (Figure 2e), the pattern changed such that phenology was earlier in the first year than the second year, suggesting a mechanism of compensation to align reproductive phenology among the growing seasons.

### 4.2 Circadian clock function correlates to multiple measures of plant phenology

Given the population differentiation in phenology, we were interested to preliminarily evaluate potential underlying genetic pathways. Using existing estimates of circadian period among the genotypes utilized in this project (McMinn *et al*., 2022), we quantified the correlation between circadian period and both size and phenology. We observed a significant relationship between circadian period and phenology (Figure 5a,b). Phenological traits expressed early in life history have previously been associated with circadian period (Suárez-López *et al*., 2001; Hayama and Coupland, 2003; Brachi *et al*., 2010). Notably, the initiation of a reproductive inflorescence from a vegetative floral rosette (bolting) has often been associated with the circadian clock (Wang and Tobin, 1998; Nozue *et al*., 2007; Chen *et al*., 2019); however, we surprisingly found no relationship between the timing of bolting and circadian period in our common garden but found strong correlations in the later expressed phenological traits (flowering, initial fruit production and maturation, and completion of reproduction). Although the initiation of bolting may be difficult to score, we are confident that any small errors will not change the overall pattern observed. We found that longer circadian periods were correlated with increased time to a given phenological stage, both among the maternal families within the BBL population and among all sampled populations. The relationship between circadian period was also expressed with the geometric mean of phenology, accounting for all three years, even though phenological timing varied across years. In *B. stricta*, the extent of linkage disequilibrium (LD) decays rapidly and is estimated as 10kb (Song *et al*., 2009), suggesting the hypothesis that the observed circadian clock-phenology correlation arises from pleiotropy of clock loci rather than LD between independently acting circadian clock and phenology genes. Although any adaptive hypothesis requires mechanistic validation, the correlation observed in the populations is consistent with the view that selection on phenology and population genetic differentiation might arise from allele frequency changes at clock loci.

The observed positive association between the circadian clock and flowering is also consistent with annotated gene function (reviewed in Greenham and McClung, 2015). In *Arabidopsis thaliana*, the control of flowering time is via the molecular pathways of the circadian clock and directly associate with the *FLOWERING LOCUS T* (*FT*) gene (Doyle *et al*., 2002; McClung, 2006; Nusinow *et al*., 2011). Both the *GIGANTEA* (*GI*) gene (Mizoguchi *et al*., 2005; Dalchau *et al*., 2011; Xie *et al*., 2015) and the clock gene *CONSTANS* (Suarez-Lopez, 2001; Valverde, 2011) are upstream of *FT* and correspond to variation in flowering time. In a separate molecular pathway, allelic variation for *COLD-REGULATED GENE 28* (*COR28*) generates long-period phenotypes with delayed flowering time and is itself regulated upstream by the core clock gene *CIRCADIAN CLOCK-ASSOCIATED1* (*CCA1*) (Rees *et al*., 2021). Circadian clock regulation of flowering time may affect the expression of later phenologic traits; that is, the timing of flowering has downstream effects on later-expressed phenological traits. For instance, the *PSEUDO-RESPONSE REGULATOR 9* (*PPR9*) gene, part of the circadian clock, regulates the *ORESARA 1* (*ORE1*) promoter which induces leaf senescence in *A. thaliana* (Rauf *et al*., 2013; Kim *et al*., 2018). Part of the *ORE1* function is controlled by the localization of GI and EARLY FLOWERING 4 (ELF4) proteins, important components of flowering time (Kim *et al*., 2020); in essence, it is flowering time control that affects the expression of senescence patterns. However, via other signaling paths, the circadian clock may directly affect phenologic traits such as fruit and seed set timing, and the expression of these stages is not simply a downstream consequence of flowering time. After pollination has occurred, the early transition to fruit set may be influenced by auxin signaling (McAtee *et al*., 2013; Fenn and Giovannoni, 2021). In *A. thaliana*, auxin is promoted by the *REVEILLE1* (*RV1*) pathway (Rawat *et al*., 2009), which is downstream of and regulated by the circadian clock. Abscisic acid (ABA) is related to fruit and seed maturation in *A. thaliana* (Karssen *et al*., 1983; Koornneef *et al*., 1989; Holdsworth *et al*., 2008), and levels of ABA are influenced by the circadian clock (Covington *et al*., 2008; Liu *et al*., 2013). Although not confirmed in *B. stricta*, together these molecular pathways suggest that genetic controls of plant phenology are downstream of the circadian clock, and changes in circadian period will directly influence not only flowering time but subsequent phenological trait expression under some growth conditions.

### 4.3 Fitness may be driven by variable selection on the circadian clock

Fitness in plant populations is determined in part by environmental variables and is expected to decrease when individuals are grown in conditions that deviate from the local environment under which their home population has evolved. Many environmental variables may covary in a coordinated manner over an elevation gradient. Nevertheless, our analyses indicate that annual precipitation is the strongest predictor of variation in phenologic timing, and precipitation has been previously shown to exert strong selective pressure in plants (Siepielski *et al*., 2017). Annual precipitation at the home sites of the plant populations would be influenced by overwinter snowfall with high snow cover at high elevations. The difference in precipitation during the coldest quarter (corresponding with the heaviest snow of the winter) from the 20 sites and the common garden site showed a negative pattern. The effect of annual precipitation in both fecundity (R = -0.59) and overall survival (R = -0.58) shows a reduction in fitness for plants coming from divergent environments into the common garden location. Reduced duration and depth of snow cover may reduce fitness in alpine populations (Wipf *et al*., 2009); even when a warming environment creates a longer growing season, frost exposure after early growth can damage budding plants and reduce fecundity. Our results suggest that the higher elevation populations, which experience increased snow cover through the winter in their homesites, have decreased fitness in the low elevation common garden. Our findings also align with results of other studies estimating fitness in experimental transplants of *B. stricta* plants. Anderson *et al*. (2015) found a similar trend of reduced fitness across home vs foreign transplant sites across an elevation gradient. Our models of precipitation for the *B. stricta* populations indicate a decrease in fitness as the difference between home site vs experimental site conditions increases. With the contribution of circadian period and its influence on phenology (short period leads to accelerated phenology at high elevation), plants from high elevation sites will flower earlier and potentially suffer meristem limitation or simply senesce earlier in the season, explaining the reduced fruit production for high elevation populations in the common garden. Based on these results, our model of fitness for these populations of *B. stricta* is

#### Circadian period → Phenology and Plant size → Survival and Fruit production

where annual precipitation is an important abiotic factor imposing selection on size and phenology, and that differences in the preceding two traits may be achieved by evolution at circadian clock loci.

We used structural equation modeling to dissect potential patterns of direct and indirect selection on the circadian clock. Fitness for populations of *B. stricta* instead were shown to have been affected by the flowering and fruiting phenology of the plants. While a lengthened circadian period was associated with delays in all phenological stages, structural equation modeling indicated that indirect selection for plant phenology via days to flowering favored a shortened clock, but first fruit initiation favored a lengthened clock period, suggesting that for *B. stricta* these life-history transitions may be more important to fitness than the first transition to reproduction at bolting. These observations suggest two possible methods to increase fitness in *B. stricta*: 1) a short circadian period resulting in early flowering, which would benefit plants from high elevations with a short growing season, and 2) a longer circadian period that delays later phenology and allows more fruit to be produced at low elevations where the growing season is lengthened. These results suggest the hypothesis that variable selection on circadian clock outputs over elevational gradients could act to preserve genetic variation and explain observed clines in the circadian clock (McMinn *et al*., 2022). In particular, populations from high elevations express shorter circadian periods, while populations from low elevation sites express longer circadian periods. Further investigation into patterns of natural selection, the biogeography of circadian clock expression, and experimental manipulations will help test these adaptive hypotheses.

## Acknowledgements

The authors would like to thank Charley Hubbard, Colten Clark, Sarah Jean O’Neill, and Katherine Traverso for their help on this project. Funding was provided by the National Science Foundation grants IOS-144571 and EPS-1755726 to C. Weinig.

